# The Impact of Methylglyoxal and SOD1 Mutation on TDP-43 Interaction in ALS Proteinopathy

**DOI:** 10.1101/2025.09.25.678586

**Authors:** Gabriela D. Ribeiro, Daniela D. Queiroz, José R. Monteiro-Neto, Gabriel F. de Souza, Paola C.S.C. Albino, Luan H. Paranhos, Tiago Fleming Outeiro, Elis Cristina Araujo Eleutherio

**Affiliations:** Institute of Chemistry, Federal University of Rio de Janeiro (UFRJ), Rio de Janeiro, Brazil. Av. Athos da Silveira Ramos, 149, Rio de Janeiro, RJ, 21941-909, Brazil; Department of Experimental Neurodegeneration, Center for Biostructural Imaging of Neurodegeneration, University Medical Center Göttingen, Göttingen, Germany; Max Planck Institute for Multidisciplinary Sciences, 37075 Göttingen, Germany; Translational and Clinical Research Institute, Faculty of Medical Sciences, Newcastle University, Framlington Place, Newcastle Upon Tyne NE2 4HH, UK; Deutsches Zentrum für Neurodegenerative Erkrankungen (DZNE), Göttingen, Germany

**Keywords:** TDP-43, SOD1, Amyotrophic lateral sclerosis, Methylglyoxal, Proteinopathy

## Abstract

**Introduction:** Proteinopathy is a key feature of amyotrophic lateral sclerosis (ALS) that causes the loss of motor neurons. Glycated SOD1 increases the levels of phospho-TDP-43, a form that aggregates in the cytosol of neurons experiencing neurodegeneration in most ALS cases.

**Objective:** Here, we evaluated whether TDP-43 interacts with SOD1 and the impact of methylglyoxal (MGO) and G93A SOD1, found in patients, on this interaction.

**Methodology:** TDP-43-SOD1 interaction was observed in H4 cells using the bimolecular fluorescence complementation (BiFC) system.

**Results:** Exposure to MGO reduced SOD1 activity and the levels of phospho-TDP-43 only in cells expressing WT SOD1. Our results showed that both WT and G93A SOD1 interact with TDP-43 in the nucleus and cytosol, with a greater proportion of cells showing cytosolic interactions between TDP-43 and the SOD1 mutant. MGO did not affect the interaction between TDP-43 and WT SOD1; however, it did lead to an increase in cytosolic inclusions at 0.4 mM MGO, a stress that resulted in a 50% reduction in cell viability. These inclusions did not colocalize with stress granules. Treatment with Cyclosporin A, an inhibitor of calcineurin (a phosphatase that dephosphorylates TDP-43), reduced the number of cells containing TDP-43 and WT SOD1 inclusions, as well as the cells showing TDP-43 and G93A SOD1 interactions in the cytosol.

**Conclusion:** Thus, we conclude that damaged SOD1, produced by MGO, or G93A mutation disrupts TDP-43 phosphorylation, altering its location within the cell and inducing its aggregation, which are important markers of ALS.

**Summary for Social Media If Published:** @laboratorio_life

Amyotrophic lateral sclerosis (ALS) is an incurable, devastating, and progressive neurodegenerative disease. It is characterized by the accumulation of misfolded proteins, such as TDP-43 and SOD1, which causes motor neuron degeneration. The relationship between SOD1 and TDP-43 remains unclear, but glycated SOD1 increases the levels of phospho-TDP-43, a form found in cytosolic inclusions within neurons undergoing neurodegeneration in most ALS cases. Our results indicate that SOD1 interacts with TDP-43 mainly in the nucleus. However, damaged SOD1, produced by methylglyoxal, or G93A SOD1, a mutant found in patients, disrupts TDP-43 phosphorylation, altering its location within the cell and inducing its aggregation, which are important markers of ALS. We therefore conclude that SOD1 plays a crucial role in the development of the disease, making it a potential target for assessing ALS risk and developing treatments.

## Introduction

Amyotrophic lateral sclerosis (ALS) is a rare neurodegenerative disease that affects upper and lower motor neurons, leading to weakness of voluntary muscles and respiratory dysfunction^1^. With rapid progression, patients have a life expectancy of about three years after symptom onset; most cases result in respiratory failure^2^. ALS is classified as familial (5 - 10%) or sporadic (90 - 95%)^1^, and proteinopathy is a hallmark of ALS, with 97% of cases presenting TAR DNA-binding protein 43 (TDP-43) dysfunction or aggregation. However, the pathological heterogeneity of ALS has been observed in mutated Cu-Zn superoxide dismutase (SOD1) ALS cases, where TDP-43 proteinopathy was not observed, and cytoplasmic aggregates have a different composition^3^.

TDP-43 is an RNA-binding protein involved in gene expression regulation^4^. Present mainly in the nucleus, where it participates in pre-mRNA splicing, transcriptional repression, and autoregulation of its gene^5^. TDP-43 can also be shuttled to the cytoplasm, where it plays important roles on mRNA stability, transport, and the formation and regulation of stress granules^4,5^. Besides cytoplasmic aggregation, mislocalization and hyperphosphorylation can also lead to ALS progression^6^.

Although twelve phosphorylation sites in TDP-43 C-terminal domain have been related to ALS, S369, S379, S403/404, and S409/410 are more consistently described in the context of the disease^7,8^. Several studies describe TDP-43 phosphorylation as a regulatory event, controlled by kinases such as casein kinases 1 and 2 (CK1/2)^9,10^, tubulin kinases 1 and 2 (TTBK1/2)^11^, and cell division cycle 7 (CDC7)^12^. Dephosphorylation is performed by calcineurin, which colocalizes with TDP-43 in human brain lysates. This phosphatase is responsible for dephosphorylating serine sites in the C-terminal domain of the protein^13^. TDP-43 phosphorylation influences splicing regulation, cellular localization^8^, condensate formation, and solubility^7^. Calcineurin also physically interacts with SOD1, enhancing its phosphatase activity^14^ and protecting against oxidative inactivation^15^.

SOD1 is an antioxidant enzyme involved in superoxide dismutation mainly in the mitochondrial intermembrane space. Present in throughout all human tissues, it is responsible for maintaining redox homeostasis along with other enzymes^16^. SOD1 plays other roles, such as to regulate the cell response against oxidative stress in the nucleus and the shift from respiration to aerobic glycolysis. Over 200 mutations have been described for SOD1^17^, with G93A being the most studied due to its association with decreased enzymatic activity, conformational instability, increased oxidative stress, and aggregation propensity^18,19^. Damages, such as glycation, can alter wild-type (WT) SOD1 conformation and promote aggregation^20,21^. Methylglyoxal (MGO), a highly reactive glycolysis byproduct, can glycates SOD1 leading to structural instability, especially at Lys122 and Lys128^22^, and has been associated with neurodegenerative diseases^23,24^. In ALS, MGO-induced glycation can impact SOD1, reducing its activity and leading to apoptosis^20^.

Here, we examined the effects of glycation in cells expressing WT or G93A SOD1. We focused on how glycation influences SOD1 activity and the C-terminal phosphorylation of TDP-43. We also investigated if TDP-43 interacts with SOD1 (WT or mutant) in living cells and how MGO affects these interactions. Finally, building on previous findings regarding phosphoTDP-43, we explored the role of TDP-43 phosphorylation in its interaction with SOD1 and its localization within the cell.

## Material and Methods

### Cell culture and MGO stress

Human neuroglioma cells (H4) were cultured in Dulbecco’s modified Eagle’s medium (DMEM) (BR30003-05 Nova Biotecnologia), supplemented with 10 % (v/v) fetal bovine serum (FBS) (10-bio500l – Nova Biotecnologia) and 1 % (v/v) penicillin/streptomycin (BR30110-01 - Nova Biotecnologia), at 37 °C in a humidified atmosphere containing 5 % CO_2_. H4 cells were challenged with four different MGO concentrations: 0.1 mM, 0.25 mM, 0.4 mM, and 0.5 mM, 24 hours after plating or after transfection, and different experiments performed 24 hours after MGO treatment.

### MTT viability assay

After MGO challenging, the medium with MGO was changed to a DMEM medium (serum-free) with 3-(4,5-dimethylthiazol-2-yl)-2,5-diphenyl tetrazolium bromide (MTT) (100 μg/ml). After two hours, the medium with MTT was disposed of and the cells were incubated in DMSO. Cellular viability was measured using a SpectraMax M2 (Molecular Devices) at a wavelength of 570 nm^25^.

### Cell transfections

To analyze the effect of MGO on SOD1 activity and phospho-TDP-43 levels, H4 cells (5,0 x 10^5^ cells/mL) were plated in 12-well plates (Kasvi) in fresh DMEM. The transient transfections were performed using 8 μL of a 2.5 M CaCl□ solution, which was added to 62 μL of a plasmid containing the cDNA sequence of human WT *SOD1* or the mutant G93A^19^. Subsequently, 2×HBS calcium phosphate buffer [50□mM BES (N,N-bis(2-hydroxyethyl)-2-aminoethanesulfonic acid), 280□mM NaCl, 1.5□mM Na□HPO□·2H□O, pH 7.05] was added dropwise to the mixture and thoroughly vortexed. After incubation for 20 min at room temperature, in the dark, the transfection solution was added dropwise to the cultured cells while the plate was gently agitated. Bimolecular Fluorescence Complementation (BiFC) plasmids were employed to investigate the interaction between TDP-43 and SOD1. A larger N-terminal fragment of Venus (VN, corresponding to 1-158 amino acid sequence), and a smaller C-terminal fragment (VC, corresponding to 159-239 amino acid sequence) were used^19^. Human TDP-43 cDNA was cloned to the 3′-end of the VN-fragment (VN-TDP-43) and human SOD1 cDNA (WT or G93A) was cloned to upstream of the VC-fragment (SOD1-VC). Transfections were performed by calcium phosphate using equal amounts of plasmids encoding VN-TDP-43 and SOD1-VC (WT or G93A SOD1). Transfection protocols were performed 24 hours after plating.

### Immunoblotting and SOD1 activity

Forty-eight hours after transfection with WT SOD1 or G93A SOD1, cells were washed with PBS and then harvested. Cell extracts for enzymatic determinations were obtained by cell incubation in a lysis buffer [NP-40 1 % (v/v), 150 mM NaCl and 50 mM Tris-HCl buffer pH 8] containing a protease inhibition mixture (cOmplete, mini, EDTA-free protease mixture inhibitor tablets; Sigma-Aldrich). Protein concentration was determined using a Bradford assay (B6916 – Sigma-Aldrich) using BSA as a standard. The gels were loaded with 25 μg of protein extract after denaturation for 10 min at 100 ºC in protein sample buffer (125.0 mM of 1.0 M Tris HCl pH 6.8, 4 % SDS 0,5 % Bromophenol blue, 4.0 mM EDTA 20 % Glycerol 10 % b-Mercaptoethanol). The samples were separated on a 12 % (w/v) SDS-polyacrylamide gel (SDS-PAGE) with a constant voltage of 130 V using running buffer (Tris-Glycine SDS 0.5 % -25.0 mM Tris, 192 mM Glycine, pH 8.3) for 1.5 hours. The samples were transferred to a nitrocellulose membrane during 2 hours with constant current at 0.35 mA using Tris-Glycine transfer buffer containing methanol^19^. For SOD1 levels quantification, the membrane was blocked overnight with 3 % (w/v) BSA in 1x PBS-T (50.0 mM Tris, 150.0 mM NaCl, 0.05 % Tween, pH 7.5) and incubated with primary antibodies either 1:1000 rabbit anti-SOD1 (HPA001401 – Sigma-Aldrich) and 1:10.000 rat anti-α-tubulin (MA180017 – Sigma-Aldrich) for 1 hour at room temperature. After washing step (three times) in PBS-T for 10 min, the membranes were incubated for 1 hour with secondary antibodies, at 1:10.000 anti-rabbit (Sigma-Aldrich), and at 1:10.000 anti-rat (Sigma-Aldrich) IgG conjugated with horseradish peroxidase in 3 % BSA/PBS-T. The quantification of TDP-43 and phosphorylated TDP-43 (Ser409) was performed using the method previously described and the membrane was incubated in primary antibodies either 1:1000 rabbit anti-TDP-43 (T1705 – Sigma-Aldrich) or 1:1000 rabbit anti-phosphoTDP-43 (Ser409) (SAB4200223 – Sigma-Aldrich) and 1:10.000 mouse anti-GAPDH (AM4300 – Termo Fisher Scientific) for 1 hour in 3 % BSA/PBS-T and further for 1 hour with secondary antibodies, 1:10.000 anti-rabbit and anti-mouse (Sigma-Aldrich) IgG conjugated with horseradish peroxidase. Detection was provided by ECL western blotting substrate (Promega) and SOD1, TDP-43 and phosphoTDP-43 levels were analyzed by ImageJ software. SOD1 activity was measured *in situ* after native polyacrylamide gel electrophoresis from 25 μg of protein in the presence of nitroblue tetrazolium (NBT) and riboflavin. The activity was determined based on the ability of superoxide dismutase to inhibit the reduction of NBT by superoxide radical. The native gel was digitized on the EC3 imaging system and SOD1 bands were analyzed using Image J software, taking into consideration the area density of all SOD1 bands^26^. Activity was expressed as the ratio between SOD1 activity and SOD1 protein level.

### Fluorescence microscopy

H4 cells transfected with VN-TDP-43 and VC-SOD1 plasmids (WT or G93A) ^19^ were stressed with MGO. Forty-eight hours after transfection with VN-TDP-43 and SOD1-VC (WT or G93A mutant), H4 cells were fixed with 4 % paraformaldehyde for 30 min at room temperature and washed three times with PBS. For nucleus stain, cells were incubated in DAPI (0.2 μg/ml) for 10 min and washed three times with PBS before the slide preparation. Fluorescence images were captured with an Olympus IX73 fluorescence microscope, and the fluorescent cells were counted. Cells were classified in two groups: cells with nuclear localization or cytoplasmic localization. Additionally, cells with TDP-43-SOD1 inclusions were counted and classified according to the total number of fluorescent cells. Results reflect the counting of at least 100 fluorescent cells per condition.

### TDP-43-SOD1 colocalization with stress granules

H4 cells treated with MGO were submitted to the immunocytochemistry (ICC) protocol 48 hours after the transfection with VN-TDP-43 and SOD1-VC plasmids. Cells were washed with PBS and fixed with 4 % paraformaldehyde for 30 min at room temperature. After fixation, cells were permeabilized with PBS 0.5 % Triton X-100 for 15 min and blocked for 1 hour with 3 % BSA in PBS-T at room temperature, followed by incubation, overnight at 4 ºC, with primary antibody 1:200 anti-G3BP1 (SAB5702394 – Sigma-Aldrich), which marks stress granules. Cells were washed three times with PBS-T and incubated with secondary antibody 1:2000 anti-rabbit IgG conjugated with Alexa Fluor 594 (A11012 – Invitrogen, ThermoFisher Scientific) for 2 hours at room temperature. Nucleus stain was obtained by DAPI and fluorescence images were captured with an Olympus IX73 fluorescence microscope^20^.

### Statistical analysis

Data was analyzed using GraphPad Prism 10.3 software and expressed as Mean ± SD of at least 3 independent experiments. Statistical differences were calculated using Two-Way ANOVA or One sample t test, as described in the figure subtitle. Significance was expressed: * P < 0.05, ** P < 0.01, *** P < 0.001, **** P < 0.0001.

## Results

### MGO decreases cellular viability

First, to evaluate MGO toxicity in the cell model used in our study (human H4 cells), we performed a cellular viability assay using MTT. This assay measures cell metabolism by converting MTT into a formazan product^25^. We tested four different MGO concentrations and observed that toxicity occurred in a dose-dependent manner, increasing with the concentration of MGO (Fig. 1). The results indicated that concentrations of 0.1 mM and 0.25 mM exhibited high viability, approaching 100 % and 80 %, respectively, whereas a concentration of 0.5 mM MGO resulted in significant cell mortality, leading to a viability of approximately 30 %. Consequently, we conducted the subsequent experiments using a MGO concentration of 0.4 mM, with 50 % of viable cells.

**Figure 1.**
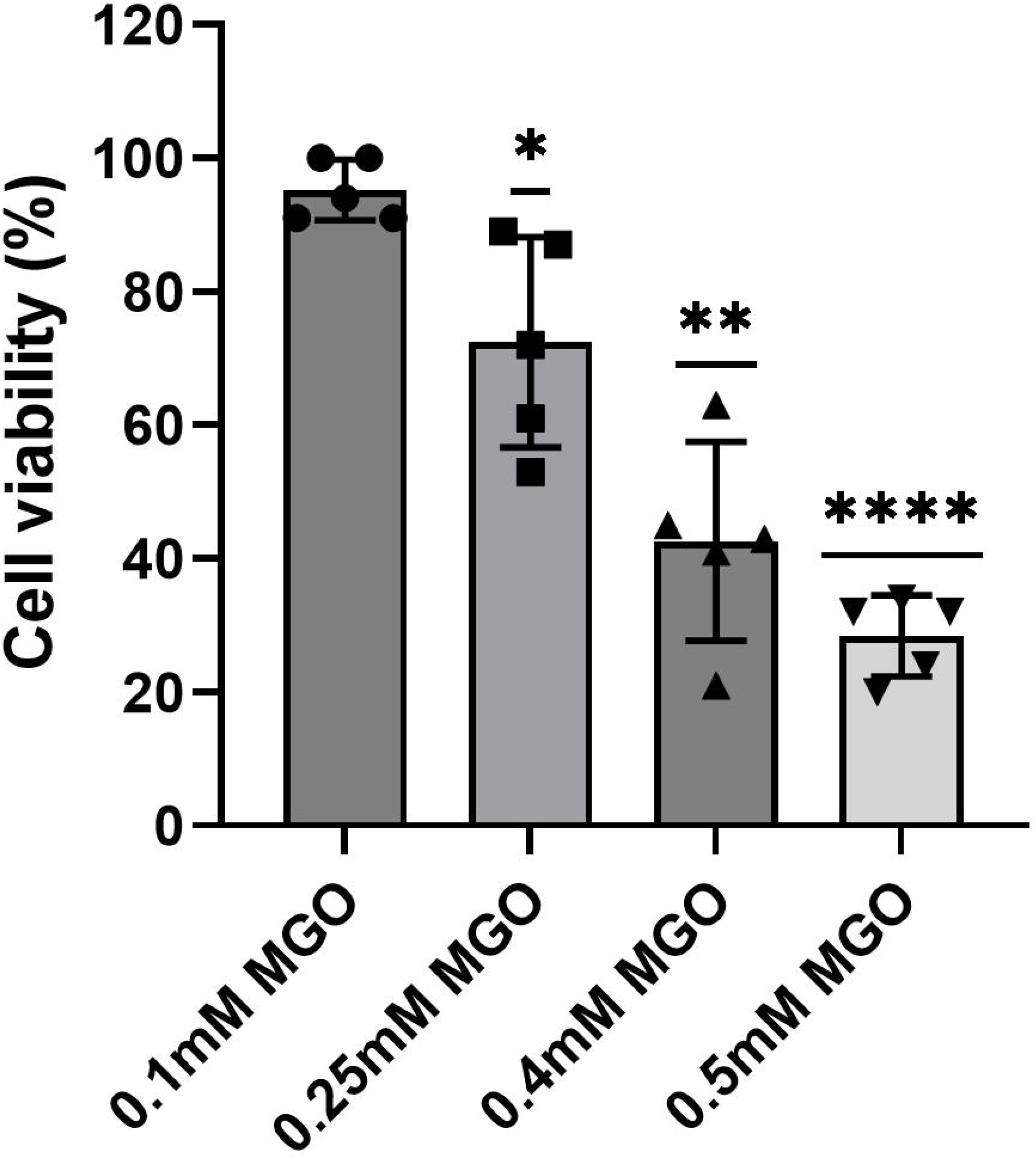
Cellular viability after MGO treatment. H4 cells were treated with four different MGO concentrations for 24 hours. The cellular viability was determinate by MTT assay and the results were expressed as Mean ± SD of the percentage of viable cells. At least three independent experiments were performed, and One-sample t test was used for statistical analysis with significance level of * p < 0.05, ** p < 0.01 and **** p < 0.0001.

### MGO stress decreases SOD1 activity and TDP-43 phosphorylation, while mutation only decreases enzyme activity

In order to assess the effect of MGO on the relative activities of WT SOD1 and G93A SOD1 mutant, we transfected H4 cells with plasmids containing the sequences of WT or G93A SOD1 and measured superoxide activity and SOD1 levels. Cells expressing mutant G93A SOD1 showed a lower SOD1 relative activity than WT SOD1 cells, even when endogenous SOD1 was expressed (Fig. 2A and S1). MGO stress decreased WT SOD1 activity (Fig. 2A), supporting previous results^20^. Interestingly, WT SOD1 activity after MGO presented levels similar to the G93A SOD1 mutant, while MGO had no significant effect on mutant activity. To assess the effect of SOD1 mutation or MGO stress on TDP-43 and on phosphoTDP-43 levels, we performed immunoblotting assays for these targets. The levels of TDP-43 were not significantly different between WT and mutant G93A cells (Fig. 2B). When we analyzed the phosphoTDP-43 levels (Ser409), we did not observe a significantly difference of phosphorylated TDP-43 in cells expressing G93A SOD1 or WT SOD1 (Fig. 2C). The phosphorylation levels decreased in WT SOD1 cells under MGO stress; however, MGO did not significantly affect TDP-43 phosphorylation in G93A SOD1 cells (Fig. 2C). Relative TDP-43 phosphorylation (Ser409) by TDP-43 levels did not demonstrate alterations on phosphorylation in cells expressing G93A SOD1, while the phosphorylation levels in WT SOD1 cells decreased with MGO stress (Fig. 2D).

**Figure 2.**
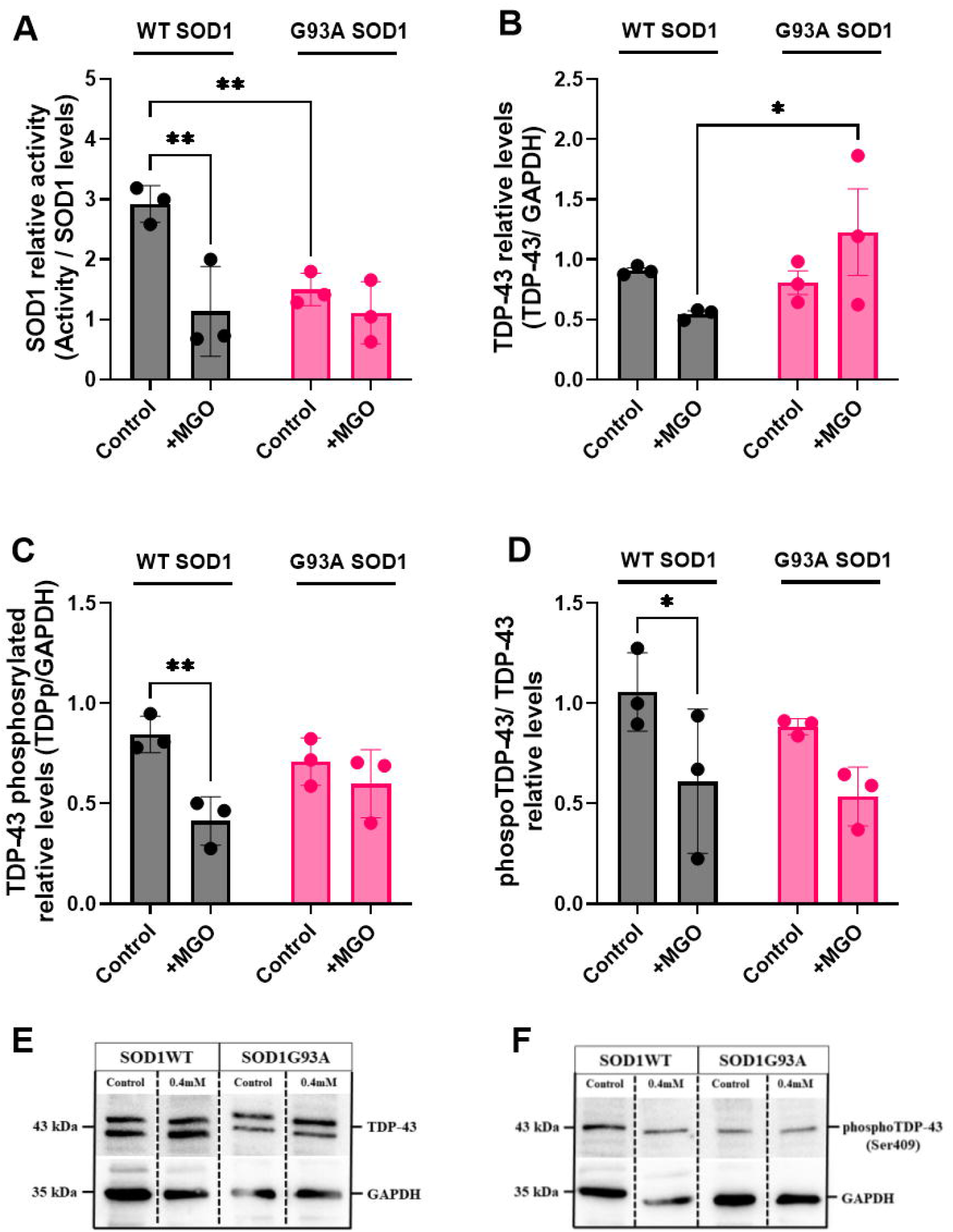
Decrease in SOD1 activity for WT and mutant SOD1 and decrease in PhosphoTDP-43 in cells expressing WT SOD1 after an MGO treatment. Cells expressing WT SOD1 or G93A SOD1 was treated with 0.4mM of MGO for 24 hours and (A) SOD1 activities were measured with non-denaturing gel zymograms by using NBT staining and normalized by SOD1 protein levels. (B) TDP-43 levels and (C) phosphoTDP-43 levels was determinate by immunoblotting assay and normalized by GAPDH levels, as well as (D) the phosphoTDP-43 levels normalized by TDP-43 levels. The results were expressed as Mean ± SD of at least three independent experiments. Two-Way ANOVA was used for statistical analysis with significance level of * p < 0.05 and ** p < 0.01.

### WT SOD1 and mutant G93A SOD1 interact with TDP-43 in different cellular compartments

Next, we used the BiFC assay to analyze if TDP-43 interacts with SOD1. This assay relies on the interaction of at least two proteins for fluorescence to occur since each protein is fused to a non-fluorescent fragment of a fluorescent protein. Therefore, the BiFC assay allows us to study heterodimeric protein species, which can then assemble into larger oligomers. Here, H4 cells were transfected with plasmids containing the sequence of TDP-43 or SOD1 (WT or mutant G93A) fused each one with the sequence of the complementary fragments of the Venus fluorescent protein. Thus, the TDP-43 and SOD1 interaction restores the Venus conformation and fluorescence, allowing visualization by microscopy.

The results showed interactions between TDP-43 and either WT or G93A mutant SOD1 (Fig. 3). Furthermore, we evaluated the cellular compartment where TDP-43 and SOD1 interact and how MGO treatment might influence these interactions. As TDP-43, SOD1 can be found in cytosol and nucleus^16^. TDP-43 interaction with WT or mutant SOD1 was primarily located in the nucleus, with approximately 40 % of cells showing interactions exclusively in that compartment. However, a higher percentage of cells showing TDP-43-G93A SOD1 than TDP-43-WT SOD1 interaction in the cytosol was observed (almost 3-time higher) (Fig. 3). It could be related to the higher proportion of the mutant SOD1 in cytosol compared to the WT form^19^. MGO stress did not change the distribution of TDP-43-SOD1 interaction, except for the mutant. The exposure to MGO reduced the percentage of cells showing TDP-43-G93A interaction in cytosol, which became similar to cytosolic interactions with the WT SOD1.

**Figure 3.**
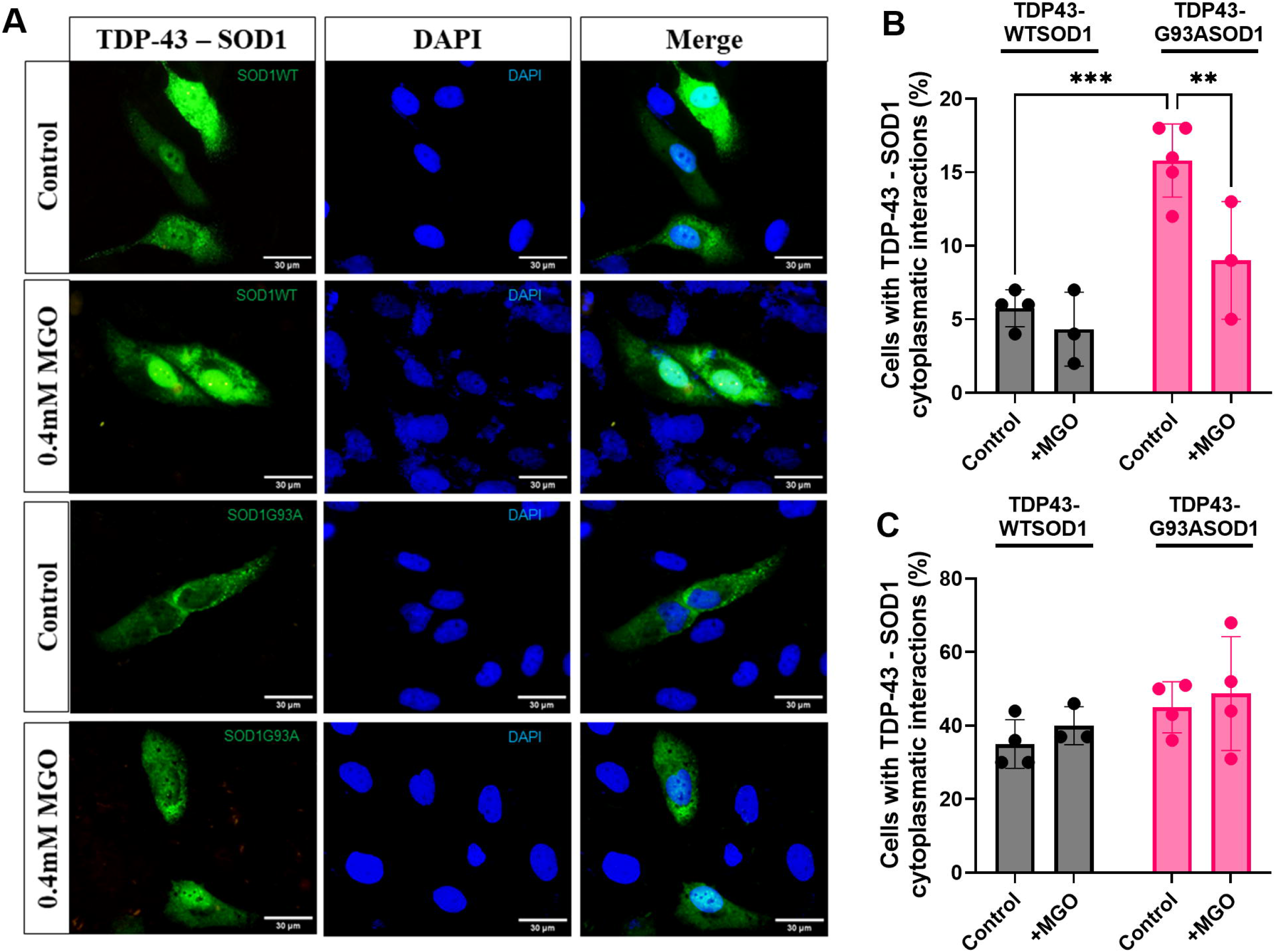
TDP-43 -SOD1G93A interactions localized in cytosol while TDP-43 – WT SOD1 localized in nuclear compartment. (A) Representative image of TDP-43-SOD1BiFC (WT SOD1 or G93A SOD1) interaction localization after 24 hours of MGO treatment. Cells were analyzed by fluorescence microscopy and fluorescents cells were counted. The results were expressed by (B) percentage of cells with TDP-43 – SOD1 interactions restricted in nucleus or (C) cytosol. The results were expressed as Mean ± SD of at least three independent experiments were performed. Two-Way ANOVA was used for statistical analysis with significance level of ** p < 0.01 and *** p < 0.001.

According to the literature, SOD1 stabilizes both CK1/2^27^, a putative kinase that phosphorylates TDP-43, and calcineurin, the phosphatase which dephosphorylates TDP-43 on Ser409/410^14^. However, the stabilization of SOD1 upon the CK1/2 seems to be dependent on its superoxide dismutase activity. To investigate how TDP-43 phosphorylation affects the compartment in which TDP–43–SOD1 interactions occur, H4 cells were treated with the calcineurin inhibitor cyclosporin A (CsA). According to the literature, the treatment of HEK293 cells with an inhibitor of this phosphatase resulted in accumulation of phospho-Ser 409/410-TDP 43^13^. The results revealed that nuclear TDP-43-SOD1WT interaction was favored by calcineurin inhibition, without effect on the interaction with G93A SOD1 in the nucleus (Fig. 4). At same time, calcineurin inhibition decreased the interactions in the cytosol, between both TDP-43-WT and mutant SOD1. The effect of MGO and CsA was always sensed in the cytosolic interaction in the case of the G93A SOD1 probably due to its lower proportion in the nucleus^19^.

**Figure 4.**
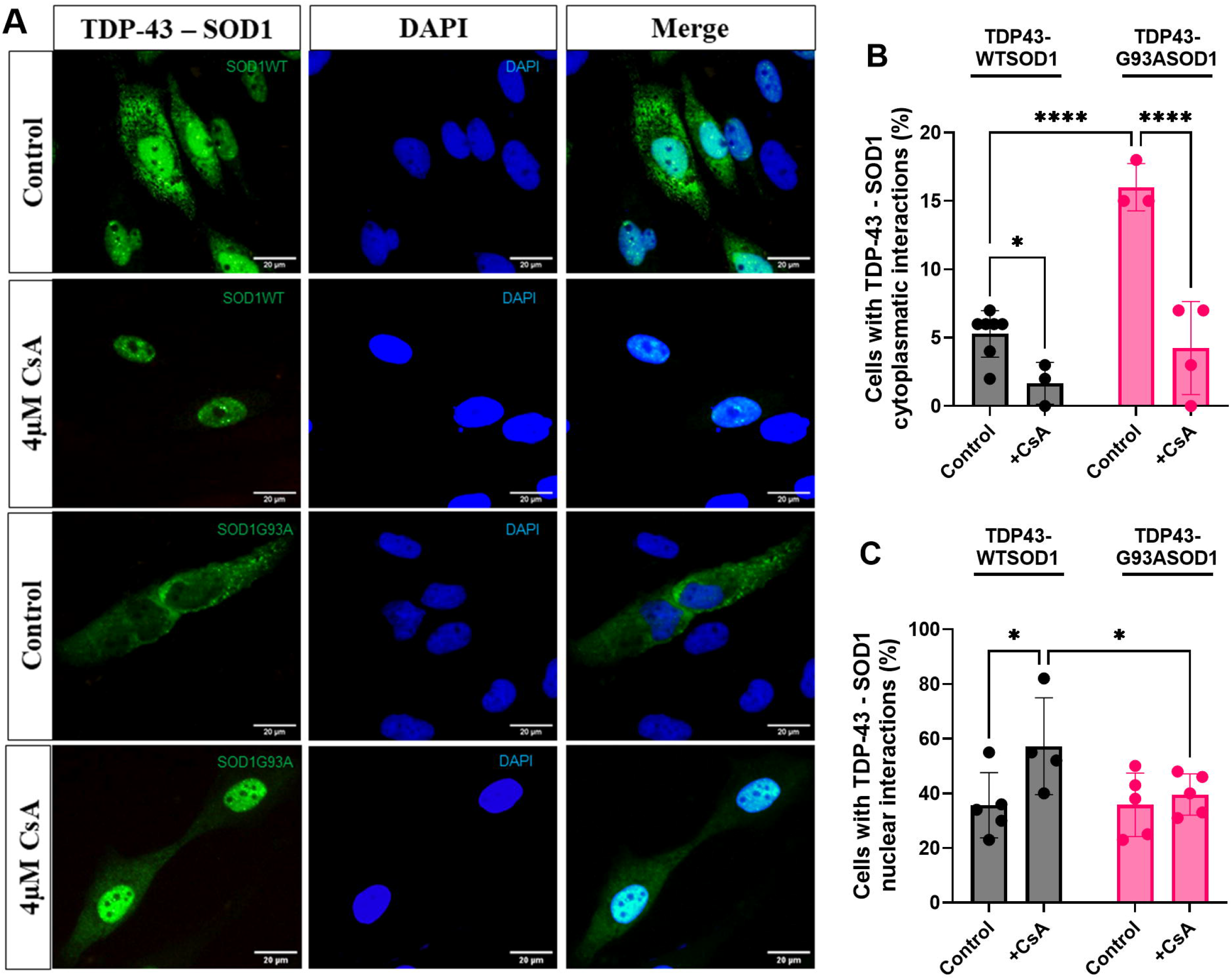
CsA treatment changed the cellular compartment where TDP-43 – WT SOD1 interactions occurs. (A) Representative image of TDP-43-SOD1BiFC (WT SOD1 or G93A SOD1) interaction localization after 24 hours of CsA treatment. Cells were analyzed by fluorescence microscopy and fluorescents cells were counted. The results were expressed by(B) percentage of cells with TDP-43 – SOD1 interactions restricted in nucleus or (C) cytosol. The results were expressed as Mean ± SD of at least three independent experiments were performed. Two-Way ANOVA was used for statistical analysis with significance level of * p< 0.05 and **** p < 0.0001.

### Methylglyoxal and calcineurin inhibition affect TDP-43 – WT SOD1 inclusion formation

Cytosolic inclusions of TDP-43 interacting with SOD1 were observed in H4 cells (Fig. 5). The inclusions were even observed in the control condition, which increased significantly in response to MGO. According to our results, MGO reduced the activity of WT SOD1 (Fig. 2A). The proportion of cells showing cytosolic inclusions of TDP-43 linked to WT SOD1 increased 2-fold after MGO stress. MGO also increased the percentage of cells with inclusions of TDP-43-G93A SOD1, however this percentage was yet lower than TDP43-WT SOD1 under control condition. Inhibition of calcineurin by CsA did not affect the proportion of cells with TDP-43-G93A SOD1 inclusions and reduced the inclusion formation with WT SOD1, although the reduction was modest (around 30 %). The main effect produced by CsA was to change the localization where TDP-43 was interacting with SOD1 (Fig. 4).

**Figure 5.**
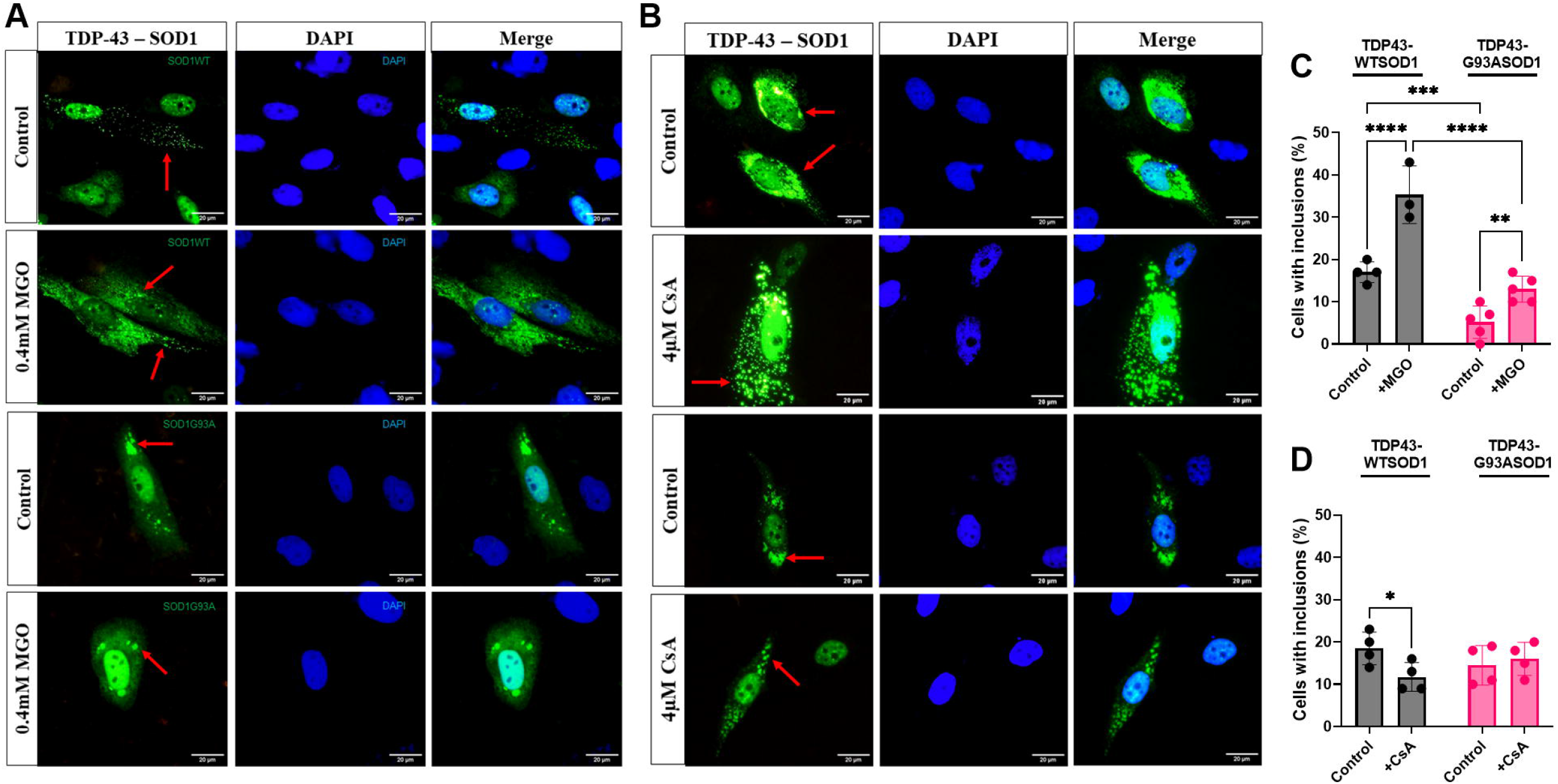
PhosphoTDP-43 decreased the TDP-43 – WT SOD1 inclusions, however did not affect the TDP-43 – SOD1G93A inclusions levels. Representative image of TDP-43-WTSOD1 or TDP-43-G93ASOD1 inclusions after 24 hours of MGO (A) or CsA (B) treatment. Cells were analyzed by fluorescence microscopy and fluorescents cells with inclusions (red arrows) were counted. The results were expressed by (B) percentage of cells with TDP-43 – SOD1 inclusions after MGO (C) or CsA (D) treatment. The results were expressed as Mean ± SD of at least three independent experiments. Two-Way ANOVA was used for statistical analysis with significance level of * p < 0.05, ** p < 0.01, *** p < 0.001 and **** p < 0.0001.

Since TDP-43 as well as glycated and mutated SOD1 can be localized in stress granules (SGs)^20^, the colocalization of TDP-43-SOD1 inclusions with SGs was evaluated. SGs were marked with G3BP1 antibody. Microscopy images showed that MGO formed cytosolic TDP-43-SOD1 inclusions; however, MGO stress was unable to form SGs, and consequently, these inclusions did not colocalize with SGs (Fig. 6).

**Figure 6.**
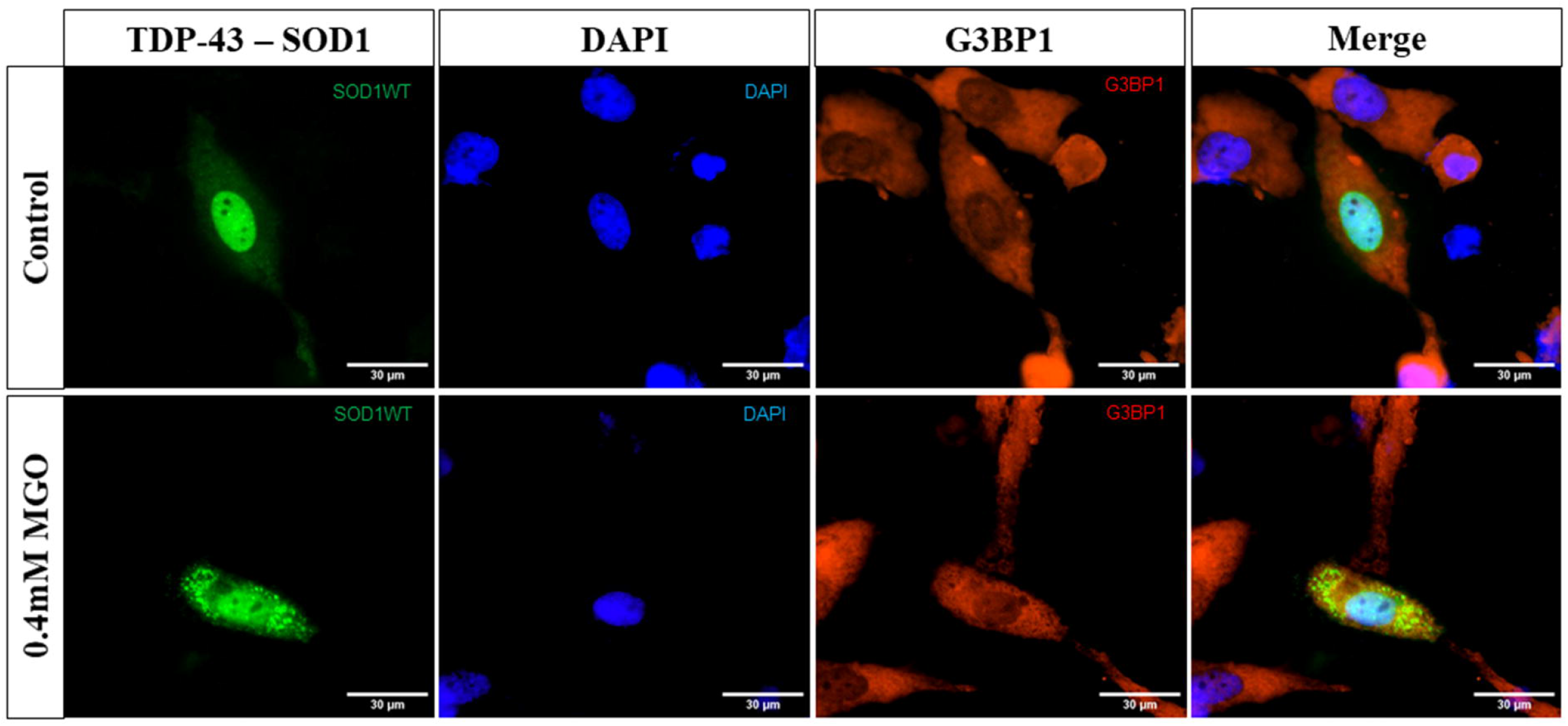
TDP-43 – WT SOD1 inclusions do not colocalize with SGs. Representative image of H4 cells expressing TDP-43-WT SOD1 after an MGO treatment. G3BP1 (red) was stained by immunocytochemistry as a marker by SGs formation and cellular nucleus was stained by DAPI (blue). Cells were analyzed by fluorescence microscopy and inclusions were observed by BiFC tagged proteins TDP-43-SOD1 (green).

Our results point out that the formation of TDP-43-WT SOD1 inclusions is correlated with phospho-TDP-43. Lower levels of phosphorylated TDP-43, as observed in response to MGO stress, increased the proportion of cells showing cytosolic inclusions of TDP-43 linked to WT SOD1. MGO impaired WT SOD1 activity which might inactivate CK1/2, impacting TDP-43 phosphorylation. On the other hand, phosphatase activity inhibition reduced the percentage of cells showing cytosolic inclusions of TDP-43-WT SOD1, although the key impact of CsA seems to be to increase the proportion of TDP-43 interacting with WT SOD1 in the nucleus.

## Discussion

This study investigated the impact of MGO stress on TDP-43 homeostasis and the interactions between TDP-43 and SOD1 in living cells. C-terminal phosphorylation of TDP-43 is a characteristic feature of ALS^7^. Calcineurin and CK1 seem to affect the amount of phosphorylated TDP-43^9,10,13^ and SOD1 might act as an intermediary, facilitating the interaction of both CK1 and calcineurin with TDP-43 to regulate the level of phosphorylated TDP-43. SOD1 binds to CK1, which is crucial for CK1’s stability; SOD1 needs to be active to prevent CK1 inactivation^27^. Conversely, SOD1 is also important for calcineurin activity. The literature has shown that calcineurin physically interacts with SOD1, boosting its phosphatase activity and stability. This interaction has been seen in the cytoplasm of neurons^14,28^.

Previous work had shown that MGO affects WT SOD1, confirming its toxic and aggregation-promoting properties^20^, as also observed in other neurodegenerative diseases ^23,29,30^. According to our results, MGO decreased viability of H4 cells, causing 50 % and 70 % mortality at concentrations of 0.4 mM and 0.5 mM, respectively. We also found that SOD1 activity significantly decreased after MGO exposure, which supports earlier findings^20^. As expected, the G93A mutation reduced SOD1 activity^31^, and MGO did not affect the activity of the mutant SOD1.

Due to the reactive nature of MGO in cells, detoxification relies on glyoxalase enzymes that convert MGO to D-lactate, in a glutathione (GSH)-dependent process. It has been observed a decrease in glyoxalase 1 activity in elderly patients^32^. Additionally, analysis of the motor cortex in ALS patients has revealed lower levels of GSH^33^, which could affect how MGO levels and oxidative stress are regulated. Consequently, genetic predispositions and unhealthy lifestyle choices, which might not be critical for young people, can become risk factors for ALS. ALS is an age-related disease with largely unknown causes.

TDP-43 and mutant SOD1 interactions were previously visualized in cells using immunoprecipitation experiments^34^; however, the TDP-43–WT SOD1 interaction had not been observed before. We used the BiFC system to analyze the physical interaction between SOD1 (WT or G93A) and TDP-43 in live cells. Our results showed that TDP-43 can interact with both WT and mutant G93A SOD1.

We observed that SOD1 binds to TDP-43 primarily in the nucleus. Furthermore, CsA, a calcineurin inhibitor, favors TDP-43 and SOD1 being present in the nucleus. A study using TDP-43 with phosphomimetic modifications in the C-terminal domain indicated that phosphorylation does not affect nuclear import or RNA regulation functions^35^. Therefore, it appears that active calcineurin favors the localization of TDP-43 and SOD1 in the cytosol. Debates about the function of TDP-43 phosphorylation suggest a regulatory role in TDP-43 localization and may prevent condensation by increasing TDP-43 solubility^7,8^.

In the cytosol, TDP-43 might be phosphorylated by CK1, which depends on active SOD1 and SOD1 binding to CK1^27^. Thus, SOD1 would be a link between TDP-43 and CK1, enabling TDP-43 phosphorylation. Since SOD1 also influences phosphatase activity, we can assume that CK1 activation would inhibit phosphatase activity. The signals that would cause SOD1 to stop activating calcineurin and start activating CK1 are unknown, but stress could be the inducer of this process.

Our results showed that phosphorylation favors TDP-43 solubility, which might facilitate its transfer to the SG. A common site of post-translational modification in TDP-43, phosphorylation of serine sites, alters the protein’s charge, increasing its solubility. This modification can change condensates to a more dynamic state, reducing aggregation and favoring SG recruitment^35^. It is unknown whether TDP-43 would travel to the SG bound to SOD1; both have been localized to the SG, especially under stress conditions.

If the stress is prolonged or very severe (such as exposure to 0.4 mM MGO, which kills 50 % of cells), SOD1 can be inactivated, inactivating CK1 and decreasing TDP-43 phosphorylation. We observed an increase in cytoplasmic SOD1-TDP-43 inclusions in response to MGO stress, without altering the cellular distribution of TDP-43-SOD1. On the other hand, treatment with CsA led to a decrease in these inclusions, albeit modest, since the cells were not under stress when treated with the calcineurin inhibitor.

Noteworthy, severe stress promotes the accumulation of misfolded proteins, overloading SGs and leading to the accumulation of cytoplasmic TDP-43 and, consequently, its aggregation^36^. This may explain why, in more advanced stages of the disease, cytoplasmic inclusions of phosphorylated TDP-43 are observed.

In the case of G93A SOD1, a greater tendency for interaction with TDP-43 in the cytosol was observed, which may be related to the mutant’s difficulty in localizing to the nucleus^19^. SOD1 plays a key role in the nucleus; in this compartment, SOD1 acts as a transcription factor, regulating genes involved in the oxidative stress response and protecting DNA from oxidative damage^37^. Therefore, the decreased activity of mutant G93A SOD1, combined with its reduced presence in the nucleus, may render the cell more susceptible to oxidative stress and DNA damage. Patients with familial SOD1-ALS present SOD1 aggregates, not TDP-43, and manifest symptoms earlier than patients with sporadic ALS. The BiFC strategy, whose fluorescence appearance is associated with the TDP-43-SOD1 interaction, did not reveal mutant SOD1 inclusions, only SOD1-linked TDP-43.

In Fig. 7, we propose a mechanism by which MGO affects TDP-43-SOD1 localization and the formation of inclusions. In conclusion, our study demonstrates the importance of SOD1 in cellular regulation and TDP-43 homeostasis. This is corroborated by Trist et al.^24^, who observed misfolded SOD1 in damaged areas showing proteinopathy of post-mortem samples from patients with sporadic and familial ALS, whether linked to SOD1 mutations or not. According to our results, a decrease in SOD1 activity, caused by MGO or mutation, triggers cellular dysregulation, leading to pro-aggregative behavior in the WT SOD1 case or mislocalization in the G93A SOD1 case. TDP-43 phosphorylation plays a crucial role in preventing aggregation, indicating increased TDP-43 solubility in WT SOD1 cells. These findings suggest that TDP-43 phosphorylation serves as a signal for nuclear localization in WT SOD1 cells. Conversely, this signaling behavior is not observed in G93A SOD1 cells, suggesting an important role for SOD1 in this regulatory mechanism.

**Figure 7.**
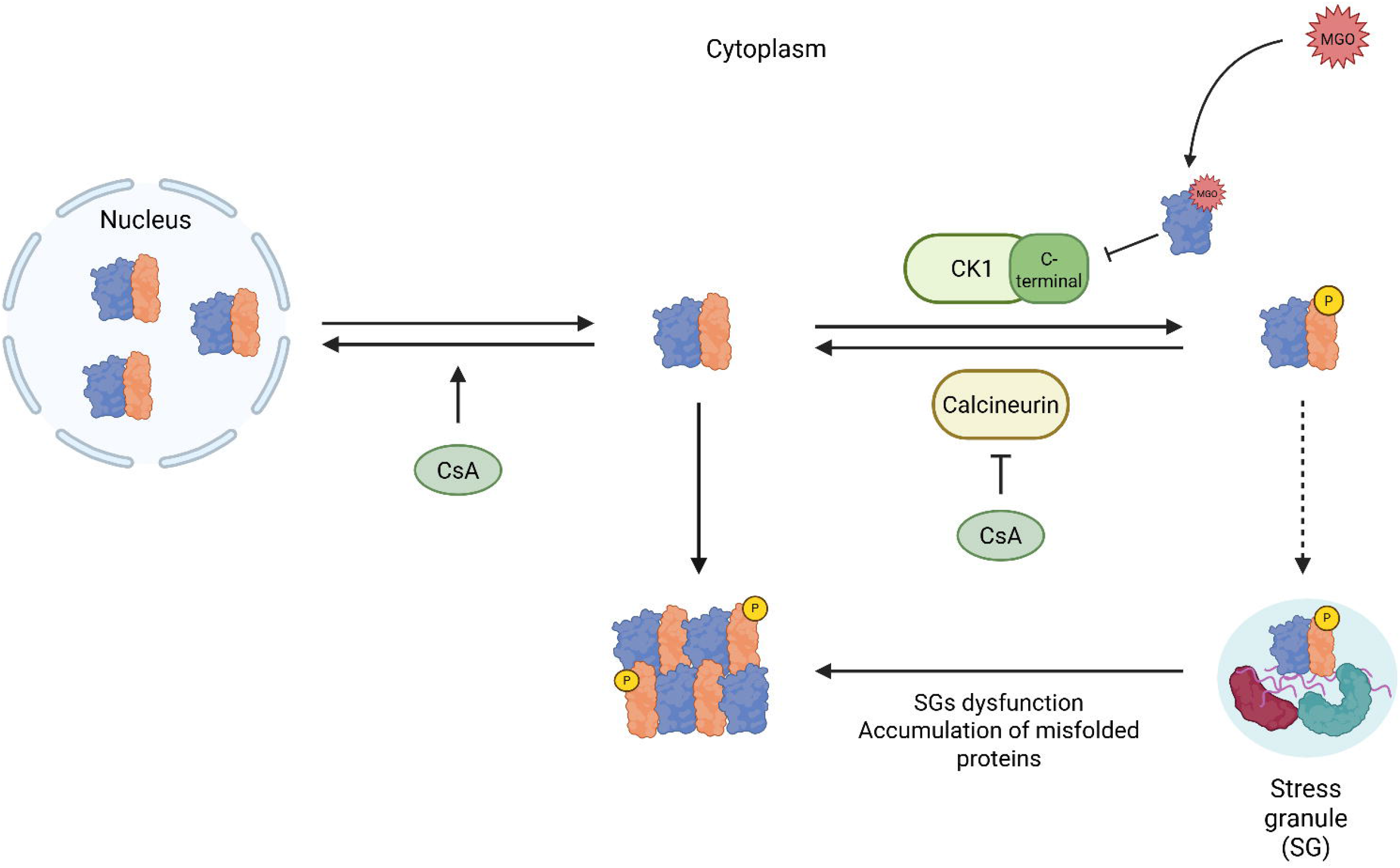
Proposed mechanism for the effect of MGO on TDP-43 – SOD1 interaction. SOD1 (blue) interacts with TDP-43 (orange) mainly in the nucleus. The level of phospho-TDP-43 (orange protein with yellow circle) appears to depend on SOD1, which influences both calcineurin and CK1. In response to MGO, SOD1 activity decreases, reducing phospho-TDP-43, which promotes inclusion formation. Phosphorylation of TDP-43 increases its solubility, reducing TDP-43 proteinopathy, a hallmark of ALS. The localization of TDP-43 in stress granules (SGs) also protects against proteinopathy. However, when the capacity of SGs to store TDP-43 is exceeded, potentially due to the accumulation of misfolded proteins within SGs, TDP-43 aggregates in the cytosol. Created in BioRender. Monteiro Neto, J. (2025) https://BioRender.com/xk4t699.

## Acknowledgements

This work was supported by grants from FAPERJ (CNE 201.174/2022), CAPES-DAAD (PROBRAL 88881.986154/2024-01), CNPq PQ (309635/2023-3) and CNPq Universal (401780/2023–6).

## Availability of data

The data that support the findings of this study are available from the corresponding author upon reasonable request.

## Author Contributions

**Gabriela D. Ribeiro –** Experimental design, Methodology, Analysis and Manuscript Writing, **Daniela D. Queiroz –** Analysis, Results Discussion, Manuscript Writing and Review, **José Raphael Monteiro Neto –** Results Discussion, Figure designs and Manuscript Writing and Review, **Gabriel F. de Souza, Paola C. S. C. Albino and Luan H. Paranhos –** Western blot analysis and Results Discussion, **Tiago F. Outeiro –** Manuscript Review, Results discussion and Funding Acquisition and **Elis C. A. Eleutherio –** Project administration, Supervision, Manuscript review, Results discussion and Funding Acquisition.

## References

1. Feldman EL, Goutman SA, Petri S, et al. Amyotrophic lateral sclerosis. The Lancet 2022;400(10360):1363–1380.

2. Masrori P, Van Damme P. Amyotrophic lateral sclerosis: a clinical review. Eur J Neurol 2020;27(10):1918–1929.

3. Mead RJ, Shan N, Reiser HJ, et al. Amyotrophic lateral sclerosis: a neurodegenerative disorder poised for successful therapeutic translation. Nat Rev Drug Discov 2023;22(3):185–212.

4. Tsekrekou M, Giannakou M, Papanikolopoulou K, Skretas G. Protein aggregation and therapeutic strategies in SOD1- and TDP-43-linked ALS. Front Mol Biosci 2024;11

5. Nilaver BI, Urbanski HF. Mechanisms underlying TDP-43 pathology and neurodegeneration: An updated Mini-Review. Front Aging Neurosci 2023;15

6. de Boer EMJ, Orie VK, Williams T, et al. TDP-43 proteinopathies: a new wave of neurodegenerative diseases. J Neurol Neurosurg Psychiatry 2021;92(1):86–95.

7. Zippo E, Dormann D, Speck T, Stelzl LS. Molecular simulations of enzymatic phosphorylation of disordered proteins and their condensates. Nat Commun 2025;16(1):4649.

8. Eck RJ, Kraemer BC, Liachko NF. Regulation of TDP-43 phosphorylation in aging and disease. Geroscience 2021;43(4):1605–1614.

9. Hasegawa M, Arai T, Nonaka T, et al. Phosphorylated TDPL43 in frontotemporal lobar degeneration and amyotrophic lateral sclerosis. Ann Neurol 2008;64(1):60–70.

10. Kametani F, Nonaka T, Suzuki T, et al. Identification of casein kinase-1 phosphorylation sites on TDP-43. Biochem Biophys Res Commun 2009;382(2):405–409.

11. Liachko NF, McMillan PJ, Strovas TJ, et al. The Tau Tubulin Kinases TTBK1/2 Promote Accumulation of Pathological TDP-43. PLoS Genet 2014;10(12):e1004803.

12. Liachko NF, McMillan PJ, Guthrie CR, et al. CDC7 inhibition blocks pathological TDPL43 phosphorylation and neurodegeneration. Ann Neurol 2013;74(1):39–52.

13. Liachko NF, Saxton AD, McMillan PJ, et al. The phosphatase calcineurin regulates pathological TDP-43 phosphorylation. Acta Neuropathol 2016;132(4)

14. Agbas A, Hui D, Wang X, et al. Activation of brain calcineurin (Cn) by Cu-Zn superoxide dismutase (SOD1) depends on direct SOD1-Cn protein interactions occurring in vitro and in vivo. Biochemical Journal 2007;405(1)

15. Ferri A, Gabbianelli R, Casciati A, et al. Oxidative inactivation of calcineurin by Cu,Zn superoxide dismutase G93A, a mutant typical of familial amyotrophic lateral sclerosis. J Neurochem 2001;79(3):531–538.

16. Eleutherio ECA, Magalhães RSS, Brasil AA, et al. SOD1, more than just an antioxidant. Arch Biochem Biophys 2021;697:108701.

17. Benatar M, Robertson J, Andersen PM. Amyotrophic lateral sclerosis caused by SOD1 variants: from genetic discovery to disease prevention. Lancet Neurol 2025;24(1):77–86.

18. Monteiro Neto JR, de Souza GF, dos Santos VM, et al. SOD1, A Crucial Protein for Neural Biochemistry: Dysfunction and Risk of Amyotrophic Lateral Sclerosis. Mol Neurobiol 2025;

19. Brasil AA, Magalhães RSS, De Carvalho MDC, et al. Implications of fALS Mutations on Sod1 Function and Oligomerization in Cell Models. Mol Neurobiol 2018;55(6):5269–5281.

20. Monteiro Neto JR, Ribeiro GD, Magalhães RSS, et al. Glycation modulates superoxide dismutase 1 aggregation and toxicity in models of sporadic amyotrophic lateral sclerosis. Biochim Biophys Acta Mol Basis Dis 2023;1869(8)

21. Banks CJ, Andersen JL. Mechanisms of SOD1 regulation by post-translational modifications. Redox Biol 2019;26:101270. [cited 2022 Oct 24]

22. Sirangelo I, Vella FM, Irace G, et al. Glycation in demetalated superoxide dismutase 1 prevents amyloid aggregation and produces cytotoxic ages adducts. Front Mol Biosci 2016;3(SEP):1–12.

23. Vicente Miranda H, ElLAgnaf OMA, Outeiro TF. Glycation in Parkinson’s disease and Alzheimer’s disease. Movement Disorders 2016;31(6):782–790.

24. Trist BG, Genoud S, Roudeau S, et al. Altered SOD1 maturation and post-translational modification in amyotrophic lateral sclerosis spinal cord. Brain 2022;00:1–23.

25. Kumar P, Nagarajan A, Uchil PD. Analysis of Cell Viability by the MTT Assay. Cold Spring Harb Protoc 2018;2018(6):469–471.

26. Brasil A de A, de Carvalho MDC, Gerhardt E, et al. Characterization of the activity, aggregation, and toxicity of heterodimers of WT and ALS-associated mutant Sod1. Proceedings of the National Academy of Sciences 2019;116(51):25991–26000.

27. Reddi AR, Culotta VC. SOD1 Integrates Signals from Oxygen and Glucose to Repress Respiration. Cell 2013;152(1–2):224–235.

28. Kim JM, Billington E, Reyes A, et al. Impaired Cu–Zn Superoxide Dismutase (SOD1) and Calcineurin (Cn) Interaction in ALS: A Presumed Consequence for TDP-43 and Zinc Aggregation in Tg SOD1G93A Rodent Spinal Cord Tissue. Neurochem Res 2019;44(1):228–233.

29. Vitek MP, Bhattacharya K, Glendening JM, et al. Advanced glycation end products contribute to amyloidosis in Alzheimer disease. Proceedings of the National Academy of Sciences 1994;91(11):4766–4770.

30. Vicente Miranda H, Gomes MA, Branco-Santos J, et al. Glycation potentiates neurodegeneration in models of Huntington’s disease. Sci Rep 2016;6(1):36798.

31. Leykam L, Forsberg KME, Nordström U, et al. Specific analysis of SOD1 enzymatic activity in CSF from ALS patients with and without SOD1 mutations. Neurobiol Dis 2024;202:106718.

32. Kuhla B, Boeck K, Lüth HJ, et al. Age-dependent changes of glyoxalase I expression in human brain. Neurobiol Aging 2006;27(6):815–822.

33. Weiduschat N, Mao X, Hupf J, et al. Motor cortex glutathione deficit in ALS measured in vivo with the J-editing technique. Neurosci Lett 2014;570:102–107.

34. Higashi S, Tsuchiya Y, Araki T, et al. TDP-43 physically interacts with amyotrophic lateral sclerosis-linked mutant CuZn superoxide dismutase. Neurochem Int 2010;57(8):906–913.

35. Gruijs da Silva LA, Simonetti F, Hutten S, et al. DiseaseLlinked TDPL43 hyperphosphorylation suppresses TDPL43 condensation and aggregation. EMBO J 2022;41(8)

36. Yan X, Kuster D, Mohanty P, et al. Intra-condensate demixing of TDP-43 inside stress granules generates pathological aggregates. Cell 2025;188(15):4123–4140.e18.

37. Tsang CK wan, Liu Y, Thomas J, et al. Superoxide dismutase 1 acts as a nuclear transcription factor to regulate oxidative stress resistance. Nat Commun 2014;5:3446.

