## Supplementary figures and images for "The Impact of Methylglyoxal and SOD1 Mutation on TDP-43 Interaction in ALS Proteinopathy"

### Figure S1

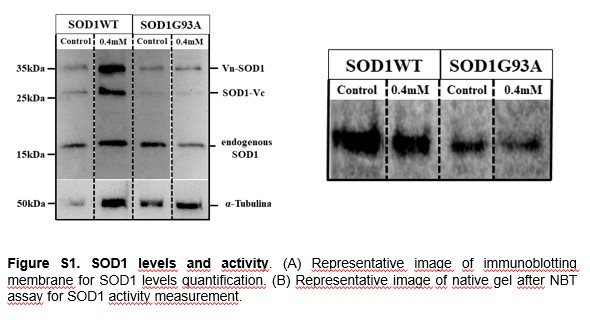
